# Aberrant hippocampal Ca^2+^ micro-waves following synapsin-dependent adeno-associated viral expression of Ca^2+^ indicators

**DOI:** 10.1101/2023.11.08.566169

**Authors:** Nicola Masala, Manuel Mittag, Eleonora Ambrad Giovannetti, Darik A. O’Neil, Fabian Distler, Peter Rupprecht, Fritjof Helmchen, Rafael Yuste, Martin Fuhrmann, Heinz Beck, Michael Wenzel, Tony Kelly

## Abstract

Genetically encoded calcium indicators (GECIs) such as GCaMP are invaluable tools in neuroscience to monitor neuronal activity using optical imaging. The viral transduction of GECIs is commonly used to target expression to specific brain regions, can be conveniently used with any mouse strain of interest without the need for prior crossing with a GECI mouse line and avoids potential hazards due to the chronic expression of GECIs during development. A key requirement for monitoring neuronal activity with an indicator is that the indicator itself minimally affects activity. Here, using common adeno-associated viral (AAV) transduction procedures, we describe spatially confined aberrant Ca^2+^ micro-waves slowly travelling through the hippocampus following expression of GCaMP6, GCaMP7 or R-CaMP1.07 driven by the synapsin promoter with AAV-dependent gene transfer, in a titre-dependent fashion. Ca^2+^ micro-waves developed in hippocampal CA1 and CA3, but not dentate gyrus (DG) nor neocortex, were typically first observed at 4 weeks after viral transduction, and persisted up to at least 8 weeks. The phenomenon was robust, observed across laboratories with various experimenters and setups. Our results indicate that aberrant hippocampal Ca^2+^ micro-waves depend on the promoter and viral titre of the GECI, density of expression as well as the targeted brain region. We used an alternative viral transduction method of GCaMP which avoids this artifact. The results show that commonly used Ca^2+^-indicator AAV transduction procedures can produce artefactual Ca^2+^ responses. Our aim is to raise awareness in the field of these artefactual transduction-induced Ca^2+^ micro-waves and we provide a potential solution.

Impact statement: Common AAV transduction procedures induce artefactual spatially confined Ca^2+^ waves in the hippocampus.

## Introduction

There has been an explosion in the use of imaging techniques to record neuronal activity over the past 30 years, starting with the introduction of organic calcium indicators to measure neuronal population activity (Yuste & Katz 1991) and accelerated by rapid advances in the development of genetically encoded Ca^2+^ indicators (GECIs) (Miyawaki *et al*. 1997). Specific advantages of neuronal Ca^2+^ imaging with GECIs lie in the ability of chronic cellular scale recordings of sizeable, densely labelled neuronal or glial populations with subtype specificity, without having to perturb the cell membrane or add synthetic chemical to the brain (Grienberger & Konnerth 2012; Rose *et al*. 2014; Semyanov *et al*. 2020).

Commonly used GECIs such as the GCaMP family have been continually improved since their initial development (Nakai *et al*. 2001), offering high signal to noise ratio, sensitivity and response kinetics such that they can detect single action potentials in vivo. This allows the reporting of cellular activity as well as the activity of sub-compartments such as the dendritic arbour (Chen *et al*. 2013; Dana *et al*. 2019; Zhang *et al*. 2023). Typically, GCaMP is expressed using transgenic animals or adeno-associated viral (AAV) transduction techniques (Tian *et al*. 2012 also see Grødem *et al*. 2023). The use of transgenic animals has the advantage of not requiring AAV transduction, thus reducing surgery load for animals and likelihood of indicator overexpression. In contrast, AAV GECI transduction is straightforward (breeding/crossing not required), can be targeted to virtually any brain region, and typically offer enhanced fluorescence (due to higher expression levels). Further, AAV transduction avoids potential hazards due to chronic GECI expression during development.

While offering unprecedented new insights into cellular scale neuronal network dynamics, it has also been reported that GECI expression in neurons can result in unwanted side-effects. Depending on the expression approach, neurons have shown reduced dendritic branching and impairment in cell health leading to cytotoxicity and cell death (Gasterstädt *et al*. 2020; Resendez *et al*. 2016). Further, increased Ca^2+^ buffering due to addition of Ca^2+^ indicators has been associated with alterations in intracellular Ca^2+^ dynamics (Grienberger & Konnerth 2012; McMahon & Jackson 2018). In addition, chronic expression of GCaMP can lead to accumulation in the nucleus and changes in gene expression (Yang *et al*. 2018). Again, depending on the specific expression approach, GCaMP variant and experimental time course, such changes may alter cellular physiology and excitability. For example, increased firing rates have been observed in hippocampal neurons expressing GCaMP5G from CaMKIIa-Cre; PC::G5-tdT mice, and epileptiform activity in neocortex in some GCaMP6 expressing transgenic mice (Gee *et al*. 2014; Steinmetz *et al*. 2017).

Here, we describe micro-scale Ca^2+^ waves that are highly confined in space, and progress slowly through the hippocampus following local GCaMP or R-CaMP viral transduction. Such aberrant hippocampal waves were typically first observed 4 weeks following injection of commercially available AAVs expressing GCaMP6, GCaMP7 or R-CaMP1.07 under the synapsin promoter. The phenomenon occurred upon GECI transduction in CA1 and CA3, but not in dentate gyrus (DG) nor neocortex, was robustly observed across laboratories with various experimenters and set-ups, and highlights the necessity of careful use of transduction methods and control measures. Reducing the transduction titre diminished the likelihood of aberrant hippocampal Ca^2+^ waves, and an alternative viral transduction method employing sparser and Cre-dependent GCaMP6s expression in principal cells avoided the aberrant Ca^2+^ waves. Further, in three transgenic GCaMP mouse lines (thy1-GCaMP6s or 6f; Vglut1-IRES2-Cre-D x Ai162(TIT2L-GC6s-ICL-tTA2), aberrant Ca^2+^ micro-waves were never observed. The aim of this article is to raise awareness in the field of artefactual transduction-induced Ca^2+^ waves and encourage others to carefully evaluate their Ca^2+^ indicator expression approach before embarking on chronic in vivo calcium imaging of the hippocampus.

## RESULTS

### Aberrant Ca^2+^ micro-wave progression through hippocampus

Based on published protocols, we injected AAV1 particles (pAAV.Syn.GCaMP6s.WPRE.SV40, Addgene #100843, titre 1×10¹³ vg/ml) in the hippocampus (total injection vol.: 500nl undiluted [1:1] virus solution) of C57BL6 wild-type animals (6 weeks old), and performed in vivo two-photon imaging to record cellular activity at 2, 4 and 6-8 weeks (wks) post-injection (p.i.) (**Fig. 1a**). Viral transduction resulted in GCaMP6s expression throughout the hippocampal CA1, CA3 and DG areas under the imaging window (**Fig. 1b**). As expected, the expression was primarily restricted to the ipsilateral hippocampus, with some labelling of projection pathways also in the contralateral hippocampus. There was no evidence of gross transduction-related morphological changes to the hippocampus, (see **Fig. 1b**) with no changes in CA1 pyramidal cell layer thickness or CA1 thickness (pyramidal layer thickness: 49 ± 12.5 µm ipsilateral and 50.3 ± 11.1 µm contralateral, n=4, Student’s t-test p=0.89; CA1 thickness: 553.3 ± 14 µm ipsilateral and 555.8 ± 62 µm contralateral, n=4, Student’s t-test p=0.94; 48 ± 13 weeks post injection at time of perfusion). At 4 wks after injection, a time point commonly used for imaging cellular activity, we observed distinctive aberrant microscale Ca^2+^ waves that travelled through CA1 recruiting neighbouring cells (**Fig. 1c-d**, **Supplementary video 1**, n=4 mice). Ca^2+^ micro-waves were maintained up to 6-8 wks after AAV injections (**Fig. 1e**, n=4 mice). In wildtype mice, these Ca^2+^ micro-waves were not observed at an earlier timepoint (2 wks p.i.; p<0.05 using Kruskal-Wallis H-test for comparison between the 3 time points).

**Figure 1:**
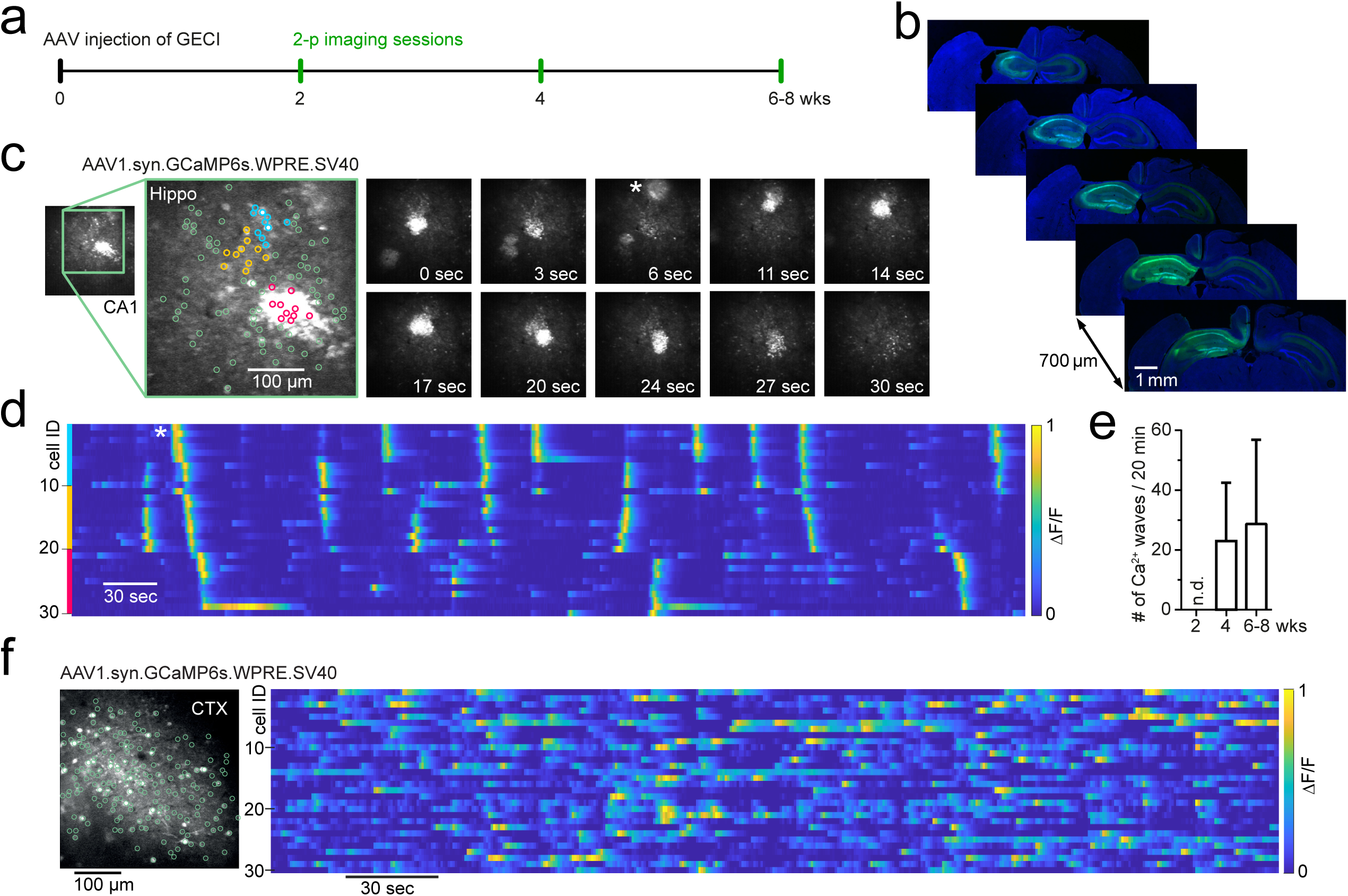
Development of Ca^2+^ micro-waves travelling through hippocampus following GCaMP transduction. **a:** Experimental protocol to examine CA1 neuronal activity using two-photon imaging following AAV transduction of genetically encoded Ca^2+^ indicators. **b:** Immunohistochemical sections following last imaging session. GCaMP6s (AAV1.syn.GCaMP6s.SV40, addgene #100843) expression throughout the ipsilateral hippocampus and projection pathways in the contralateral hippocampus. **c**: Two-photon Ca^2+^ imaging of FOV in CA1 at 4wks p.i. showing aberrant Ca^2+^ micro-waves (see also Supplementary video 1). Magnified inset shows 3 colored neuronal subgroups (blue, orange, magenta) based on their spatial vicinity from a total population of 100 identified neurons (green). *right:* time series of two-photon Ca^2+^ imaging FOVs showing two Ca^2+^ micro-waves, the first at 0 sec, the second appearing at 6 sec (asterisk). The second wave progresses through FOV over dozens of seconds. **d**: Raster-plot of individual neuronal Ca^2+^ activity (ΔF/F, 1min moving window, traces max-normalized per neuron) from neighboring subgroups (colors correspond to c). Asterisk (same as in c): a Ca^2+^ micro-wave advances through neighboring neuronal subgroups. **e**: Occurrence rate of aberrant Ca^2+^ micro-waves with increasing expression time, following viral transduction of AAV1.syn.GCaMP6s.SV40 in mature C57BL6 wild-type animals. n.d. = none detected. **f**: Two-photon Ca^2+^ imaging FOV in the visual cortex at 6wks p.i. (left) with normal sparse spontaneous Ca^2+^ activity and no detected Ca^2+^ micro-waves (right; raster plot of ΔF/F, 1min moving window, traces max-normalized per neuron).

The properties of the Ca^2+^ micro-waves depended on the hippocampal region and exact recording location. For instance, although the Ca^2+^ waves were consistently observed in CA1, the spatial dimensions of the Ca^2+^ micro-waves were broader in the stratum oriens compared with stratum pyramidale of CA1 (**Supplementary videos 4 [str. pyr.] and 5 [str. oriens]**), which likely reflects concomitant neuropil activation. We next examined, if the CA1 network is particularly prone to the generation of such waves and if they show regional specificity. Upon viral GCaMP6s transduction under synapsin, Ca^2+^ waves were observed in both CA1 (n=4/4; **Supplementary videos 1-5**) and CA3 (n=1/1; **Supplementary video 6**), but interestingly, not in the dentate gyrus (n=3 mice, 4, 8 & 10 wks p.i., 40 min total recording time per mouse). In contrast to hippocampus, synapsin dependent GCaMP6s expression restricted to the neocortex (V1 or somatosensory cortices) did not result in cortical Ca^2+^ waves in our hands (**Fig. 1f**, n = >20 mice).

### Aberrant Ca^2+^ microwaves in disease models

The observed Ca^2+^ micro-waves were distinct from local seizure activity (no rhythmicity, no typical ictal evolution, no postictal depression) (Masala *et al*. 2022; Muldoon *et al*. 2015; Wenzel *et al*. 2017, 2019a) and spreading depolarization/depression phenomena (no concentric expansion, no post-wave neural depression). However, the occurrence of these artificial events may be confused as aberrant activity related to a pathology, especially when studying pathologies with known cellular and network hyperexcitability. For example, we initially found the aberrant hippocampal Ca^2+^ micro-waves in the *Scn2a*^A263V^ model of genetic epilepsy; however, these Ca^2+^ waves in CA1 of heterozygous animals (5/5 mice) were in general similar to those detected in wildtype animals at 4 wks p.i.. In the *Scn2a*^A263V^ model, in one case (1/5 animals), Ca^2+^ waves were observed even at 2 wks p.i. (**Supplementary video 2**). Furthermore, hippocampal transduction of jGCaMP7f under synapsin (Addgene #104488, AAV9 particles, original titre 2.5×10¹³ vg/ml, total inj. vol. 1000nl [1:2 dilution]) in a mouse model of Alzheimer’s disease (PV-Cre::APPswe/PS1dE9) also resulted in Ca^2+^ micro-waves (n=3/6 mice). Together, these experiments show that common AAV injection procedures of GECIs under the synapsin promoter lead to artefactual hippocampal Ca^2+^ micro-waves in wildtype mice and genetic mouse models of disease.

### Properties & robustness of aberrant hippocampal Ca^2+^ waves

Next, we investigated the robustness of the aberrant Ca^2+^ micro-waves across institutes and conditions. We chose to compare the incidence of aberrant Ca^2+^ micro-waves in the CA1 region in four separate institutes in three different countries following transduction of GCaMP6s (Addgene #100843; IEECR/UoB, CU), GCaMP6m or jGCaMP7f (Addgene #100841 or #104488; DZNE) or RCaMP1.07 (Viral Vector Facility UZH #V224-9; UZH, **Supplementary video 3**)(**Table 1**).

**Table 1:**
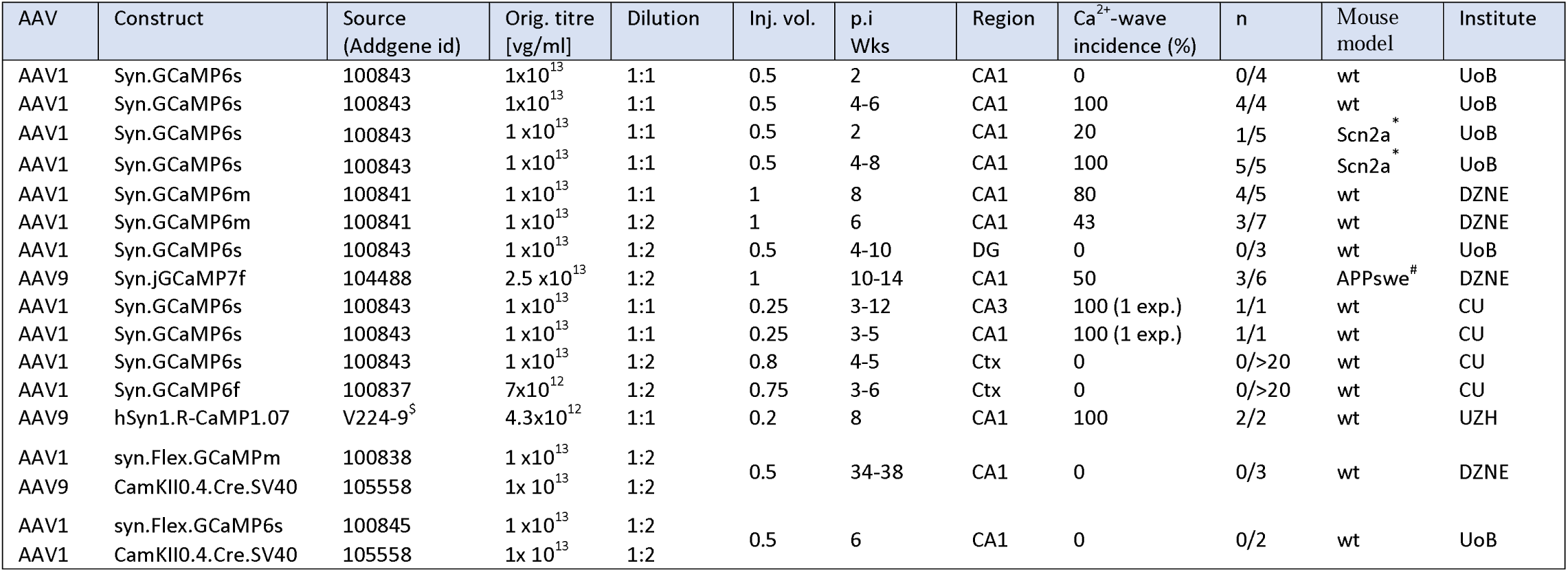
Viruses used for expression of GECI. Viral titre is from Addgene documentation and was used at original concentration (dilution of 1:1) or at a dilution of 1:2. Syn.Flex.GCaMP6s and CamKII0.4.Cre were co-injected and therefore diluted to 1:2. Two-photon Ca^2+^ imaging was performed from 2 weeks after injection in the hippocampus (CA1, CA3 or DG) or neocortex (Ctx). Ca^2+^ micro-wave incidence was determined from the number of animals exhibiting Ca^2+^ micro-waves at the specified timepoint and region. * in heterozygous *Scn2a*^A263V^ mice. # in PV-Cre::APPswe/PS1dE9 mice. ^$^ Sourced from Viral Vector Facility University of Zurich (VVF/UZH). Inj. vol. = injection volume in µl; p.i. = post-injection; wks = weeks.

The incidence of aberrant hippocampal Ca^2+^ micro-waves was robust, observed at the 4 different institutes each using variations of commonly used, published viral transduction procedures and standard two-photon Ca^2+^ imaging protocols (**Table 1** and see methods for more details). Importantly, aside from the targeted region, the viral titre was important, as halving the original AAV1.syn.GCaMP6m viral titre decreased the number of animals that developed aberrant Ca^2+^ micro-waves from 80% of animals (4/5, original titre, 1×10¹³ vg/mL) to 43% of animals (3/7, 50% reduced titre, 0.5×10^13^ vg/ml) (see **Table 1**). To statistically test the involvement of expression level, we used a generalized linear model. For injections into CA1 in the hippocampus (n=28), a multivariate logistic GLM (Ca^2+^ wave ∼ dilution + p.i. wks) found both dilution and post injection weeks were significantly related to Ca^2+^ wave incidence (model deviation above null = 7.5; dilution: z score = 2.18, p < 0.05; p.i. wks: z score = 2.22, p < 0.05).

To examine how robust the Ca^2+^ micro-waves properties were, we compared properties across the different laboratories following expression of GCaMP6m and GCaMP6s variants. The occurrence rate of the aberrant Ca^2+^ micro-waves was similar across the different institutes (**Fig. 2a**). Ca^2+^ micro-waves were spatially confined (diameter range of 200-300 µm), moved across the field of view with slow progression speeds (speed range of 5-25 µm/sec) (**Fig. 2b)**, and displayed no rhythmicity but rather plateau-like Ca^2+^ activity. No statistical differences were observed in Ca^2+^ micro-wave properties between the different institutes suggesting that these values provide a reasonable range. In addition, the Ca^2+^ micro-waves were not restricted to a single GECI variant or version, with Ca^2+^ waves observed following expression of GCaMP6m (n=4), GCaMP6s (n=5), and GCaMP7f (n=3), as well as R-CaMP1.07 (n=1)(**Fig. 2c** and **Table 1**).

**Fig. 2.**
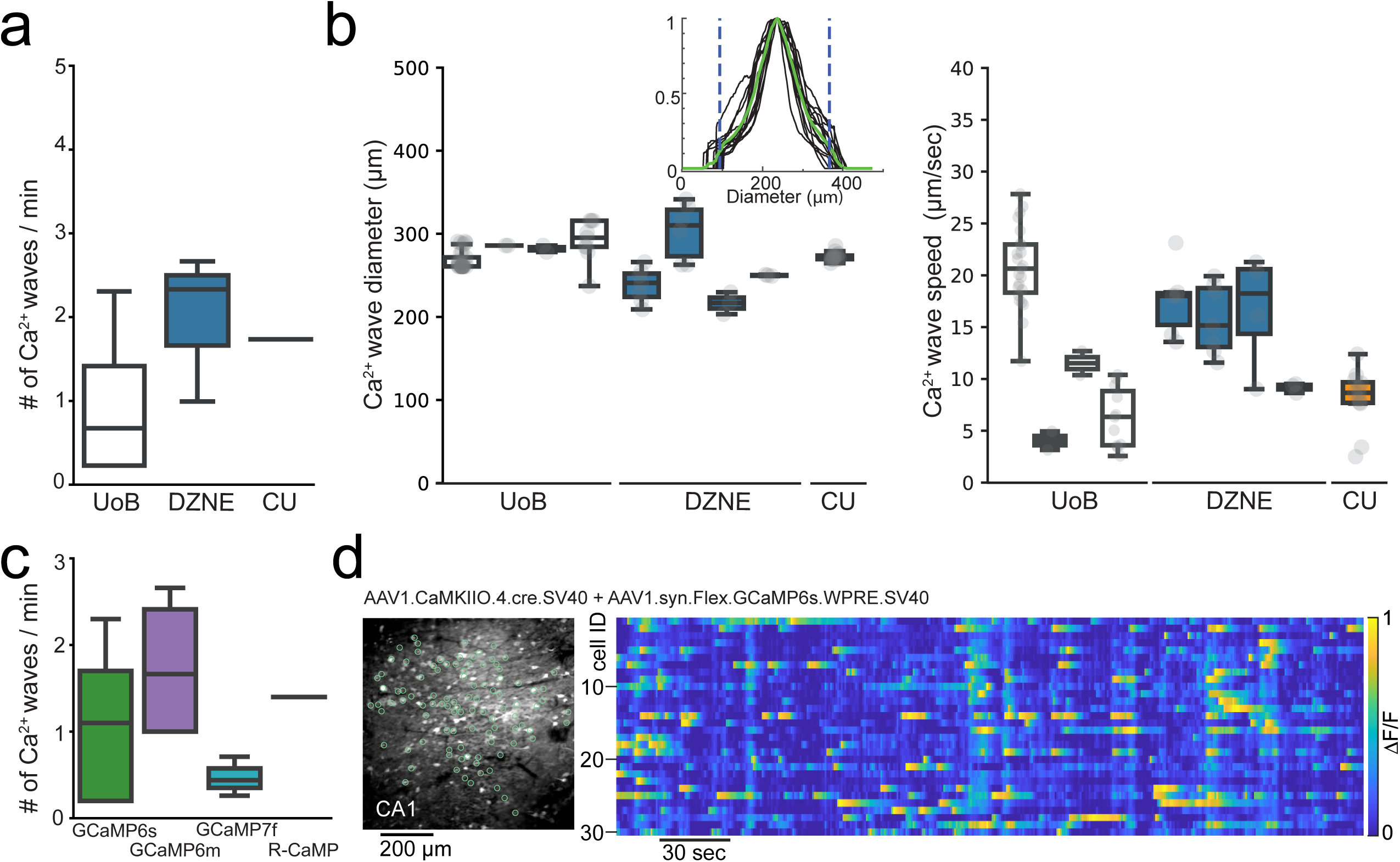
Aberrant Ca^2+^ micro-waves are consistent across laboratories and GECI variant. **a:** Plot of the occurrence rate of aberrant Ca^2+^ micro-waves in CA1 at the different institutes at 6-8wks after injection of GCaMP6s or GCaMP6m. **b:** Ca^2+^ micro-wave diameters (left) and progression speed (right) in CA1 from each animal recorded across institutes. Inset: Histogram of fluorescent intensity taken across each Ca^2+^-wave within an animal. Green line is the average, areas outside dashed lines mark 10% lowest fluorescence values, which were excluded from analysis. **c:** Plot of the occurrence rate of aberrant Ca^2+^ micro-waves in CA1 following injection with commonly used GECIs (see table 1). **d:** Two-photon Ca^2+^ imaging FOV (left) in hippocampal CA1 following dual injection approach for conditional GCaMP6s expression (6 wks p.i.) with normal sparse spontaneous Ca^2+^ activity and no detection of Ca^2+^ micro-waves (right; raster plot of ΔF/F, 1min moving window, traces max-normalized per neuron).

In summary, upon synapsin-promoter-dependent AAV Ca^2+^ indicator expression, depending on time of expression and viral transduction titre, Ca^2+^ micro-waves were specifically observed in the CA1 and CA3 subregions of the hippocampus. For CA1, the Ca^2+^ micro-waves were observed across laboratories and countries and animal models, using common transduction procedures (for overview see **Table 1**).

### Alternative transduction method of GCaMP to avoid aberrant Ca^2+^ micro-waves

In light of these results, we tested an alternative expression approach to avoid aberrant hippocampal Ca^2+^ micro-waves. To this end, we selected an approach to both limit the expression to principal cells and label a sparse population of the principal cells using a dual AAV injection approach. Here, Cre-dependent expression of GCaMP6s or GCaMP6m was achieved in a sparse population of principal cells under the CaMKII promoter (AAV1.syn.Flex.GCaMP6s.WPRE.SV40, Addgene #100845, and AAV1.CamKII0.4.Cre.SV40, Addgene #105558; n=2 or, AAV1.Syn.Flex.GCaMP6m.WPRE.SV40, Addgene #100838 and AAV9.CamKII0.4.Cre.SV40, Addgene #105558; n=3; **Fig. 2d**) (see Hare *et al*. 2022; Jimenez *et al*. 2020; Sheffield & Dombeck 2015), upon which no Ca^2+^ micro-waves were observed (0/5 animals, **Fig. 2d**). Further, hippocampal Ca^2+^ micro-waves were neither observed in transgenic thy1-GCaMP6s nor 6f mice (JAX strain 025776 or 024276; up to 3 months of chronic imaging in n>30 mature mice age >p60, cumulative imaging time >200hrs), nor in Vglut1-IRES2-Cre-D mice crossed with Ai162(TIT2L-GC6s-ICL-tTA2)-D mice (JAX strains: 037512, 031562; up to 3 months of chronic imaging in n=5 mature mice >p60).

## Discussion

Here we report titre- and expression-time-dependent aberrant hippocampal Ca^2+^ micro-waves in CA1 and CA3 regions following viral expression of GCaMP or R-CaMP1.07 under the synapsin promoter. These aberrant Ca^2+^ micro-waves robustly occurred, were observed in 4 different institutes each using a common viral transduction approach and standard two-photon Ca^2+^ imaging protocols.

Ca^2+^ micro-waves were typically first detected at ∼4wks, rarely also at 2 wks, after injection. Thus, there may be a time window when Ca^2+^ activity could be recorded in the absence of this artefactual phenomenon. However, we would still hesitate to use this specific approach for hippocampal imaging experiments as, although unknown from our data, more subtle alterations may occur prior to visible onset of aberrant activity. Further, at sites more distal to the injection site with lower expression levels, Ca^2+^ micro-waves may not be observed; however, it may very well be that Ca^2+^ micro-waves in regions with higher expression will affect fine-scaled neuronal population dynamics in primarily unaffected neighbouring regions.

The presence of Ca^2+^ micro-waves was not restricted to a single GCaMP variant or version, and was observed using either GCaMP6m, GCaMP6s, or GCaMP7f. The phenomenon was also observed upon transduction of R-CaMP1.07, indicating that these aberrant hippocampal waves are not restricted to GCaMP indicators, but rather present a general phenomenon following Ca^2+^- indicator transduction. Notably, the viral transduction titre was a key factor, as reducing the viral transduction titre from 1×10^13^ vc/ml (500nl or 1000nl of a 1:1 undiluted virus solution) to 5×10^12^ vc/ml (1000nl 1:2 solution, single injection) decreased, albeit did not yet prevent, the occurrence of Ca^2+^ micro-waves. In the literature, hippocampal GCaMP transduction procedures in mice typically include one to several separate nearby injections, with a total volume of transduced undiluted virus ranging from 60nl to 500nl (Cai *et al*. 2016; Jimenez *et al*. 2020; Keinath *et al*. 2022; Pettit *et al*. 2022; Radvansky *et al*. 2021; Skocek *et al*. 2018; Szabo *et al*. 2022; Weisenburger *et al*. 2019; Wirtshafter & Disterhoft 2022; Zaremba *et al*. 2017). In other studies, syn-GCaMP virus was diluted prior to injection (up to 1:10)(Jimenez *et al*. 2020; Zong *et al*. 2022), resulting in varied transduction volumes up to 1500nl. In our hands, a reduced viral titre of 5×10^12^ vc/ml in a 1000nl injection volume still resulted in aberrant Ca^2+^ micro-waves. Thus, viral transduction titres per volume well below this number and diluted transduction solutions are advisable for syn-GCaMP expression in the hippocampus, if AAV syn-Ca^2+^-indicator transduction is desired for a planned in vivo hippocampal imaging experiment.

If possible, alternate viral GCaMP expression approaches should be chosen. As a possible alternative, similar to previous reports using dual AAV injections or AAV in Cre-driver mouse lines (Farrell *et al*. 2020; Grosmark *et al*. 2021; Mineur *et al*. 2022; Rolotti *et al*. 2022; Terada *et al*. 2022), we find that Cre-dependent AAV GCaMP expression (IEECR/UoB) in pyramidal neurons does not cause aberrant hippocampal Ca^2+^ micro-waves. Moreover, we have not observed this aberrant phenomenon in transgenic thy1-GCaMP6s or 6f mice (JAX strain 025776 or 024276)(Masala *et al*. 2022; Rupprecht *et al*. 2023; Wenzel *et al*. 2019b), nor in Vglut1-IRES2-Cre-D x Ai162(TIT2L-GC6s-ICL-tTA2)-D mice (JAX strains: 037512, 031562). It goes beyond the scope and available resources in our laboratories to further identify, which viral GCaMP transduction approaches avoid the reported phenomenon. It seems likely that the underlying mechanisms for this artefact comprise transduction titre and time period of GCaMP or R-CaMP1.07 expression, region specificity and density of expression. Importantly, although our data suggest some regions and AAV constructs seem more prone to generate artefactual Ca^2+^ waves under certain conditions, this does not mean that Ca^2+^ waves cannot be generated in other regions, or with other constructs or promoters. It remains unclear if the observed phenomenon is restricted to Ca^2+^ indicator viral expression in mice, or if it extends to different animal models as well. In this regard, a previous report did not observe Ca^2+^-waves in rats following synapsin-dependent GCaMP6m expression, although notably, imaging was performed under isoflurane anaesthesia (Sosulina *et al*. 2021). Further, disentangling the exact cellular mechanisms of the phenomenon from technical aspects is difficult as e.g. the mere change in the transduction procedure will affect GECI expression level. For instance, although Ca^2+^ waves were not observed following conditional expression of GCaMP with CaMKII.Cre, which may suggest a requirement for interneuronal expression, it may also simply reflect differences in final GCaMP expression density and levels between the two transduction procedures.

In the context of this study, the phenomenon of Ca^2+^ micro-waves is possibly related to the expression of exogenous Ca^2+^ buffer and the resulting effects on Ca^2+^ dynamics and gene expression (McMahon & Jackson 2018; Rose *et al*. 2014; Yang *et al*. 2018), which may be why our findings extended across genetically encoded Ca^2+^ indicators. Beyond clearly being abnormal, the exact nature of the observed Ca^2+^ micro-waves remains unclear and may reflect Ca^2+^ influx during action potential firing or possibly Ca^2+^ release from internal stores. In a limited dataset, we tried to detect the Ca^2+^ micro-waves by hippocampal LFP recordings (insulated Tungsten wire, diameter ∼110µm). We could not identify a specific signature, e.g. ictal activity or LFP depression, which may correspond to these Ca^2+^ micro-waves. The shortcoming of these LFP recordings is that we could not simultaneously perform hippocampal 2-photon microscopy and thus, it is uncertain if the Ca^2+^ micro-waves indeed occurred in proximity to our electrode. We did not evaluate the effect of Ca^2+^ micro-waves on physiological activity. Based on the data presented here, it appears reasonable to hypothesize that such waves obscure if not interfere with physiological activity e.g. with hippocampal place cell activity. However, the primary purpose of this manuscript was to inform the community about an artefact that can be avoided using alternative approaches.

In summary, this report shows that common AAV hippocampal injection procedures of Ca^2+^ indicators may lead to aberrant Ca^2+^ micro-waves in wildtype mice and genetic mouse models of disease, particularly if high titre virus loads are used. The aim of this article is not to discredit Ca^2+^ indicators expressed under the synapsin promoter, a tool that we greatly appreciate ourselves, but to sensitize the field to artefactual transduction-induced aberrant Ca^2+^ micro-waves. The underlying mechanisms, some of which we have described above, are likely multi-faceted. This article seeks to inform and alert others to carefully evaluate their Ca^2+^ indicator expression approach for in vivo Ca^2+^ imaging of the hippocampus, which is becoming increasingly popular. There is certainly a much greater number of safe alternate hippocampal Ca^2+^ indicator viral expression approaches than has been reported here, and we encourage others to report on viral Ca^2+^ indicator transduction safety profiles. Indeed, others have also encountered these artefactual events as recent social media posts attest (Application Specialist Team 2023). With more indicators of brain cell activity becoming available (Ca^2+^ indicators and others including voltage indicators) as well as routes for viral delivery (Grødem *et al*. 2023), the open and timely reporting of transduction safety profiles will reduce unnecessary animal experiments, save laboratory resources and time in future investigations into hippocampal function in health and disease.

## Materials and Methods

### Animals

All experiments followed the institutional guidelines of the Animal Care and Use committee of respective institutions (University of Bonn, DZNE and Columbia university). We used wild-type C57BL/6J mice, Thy1-GCaMP6 mice (C57BL/6J-Tg(Thy1-GCaMP6s)GP4.12Dkim/J; Jackson Lab Stock No: 025776, or C57BL/6J-Tg(Thy1-GCaMP6f)GP5.5Dkim/J; Jackson Lab Stock No: 024276 (Dana *et al*. 2014)), Vglut1-IRES2-Cre-D mice (Jackson Lab Stock No: 037512) crossed with Ai162(TIT2L-GC6s-ICL-tTA2)-D mice (Jackson Lab Stock No: 031562), PV-Cre::APPswe/PS1dE9 (cross between B6;129P2-Pvalbtm1(cre)Arbr/J, Jackson Lab Stock No: 008069, and B6.Cg-Tg(APPswe,PSEN1dE9)85Dbo/Mmjax, Jackson Lab Stock No: 034832) or *Scn2a*^A263V^ mice (from Dirk Isbrandt see Schattling *et al*. 2016). Mice were kept under a light schedule of 12h on/12h off, constant temperature of 22 ± 2°C, and humidity of 65%. They had ad libitum access to water and standard laboratory food at all time. All efforts were made to minimize animal suffering and to reduce the number of animals used.

### Virus injections

For in vivo two-photon imaging experiments, GECIs were virally transduced using injection of an AAV (see **Table 1** and **Table 2**). At the time of injection mice ranged in age from 5 – 79 wks. There was no significant relationship between the age of the animal and the incidence nor frequency of Ca^2+^ micro-waves during this period (linear regression fit to the Ca^2+^ wave frequency against age was not significant: intercept = 1.37, slope = -0.007, p=0.62, n = 14; and generalized linear model relating Ca^2+^ wave incidence ∼ age was not significant: z score = 0.19, deviance above null = 0.04, p = 0.85, n=24).

**Table 2:** Key Resources.

| Construct | Full plasmid | AAV | Source | ID |
| --- | --- | --- | --- | --- |
| Syn.GCaMP6s | Syn.GCaMP6s.WPRE.SV40 | AAV1 | Addgene | 100843 |
| Syn.GCaMP6m | Syn.GCaMP6m.WPRE.SV40 | AAV1 | Addgene | 100841 |
| Syn.jGCaMP7f | Syn-jGCaMP7f-WPRE | AAV9 | Addgene | 104488 |
| Syn.GCaMP6f | Syn.GCaMP6f.WPRE.SV40 | AAV1 | Addgene | 100837 |
| hSyn1.R-CaMP1.07 | hSyn1-chI-RCaMP1.07-WPRE-SV40p(A) | AAV9 | VVF/UZH | V224-9 |
| syn.Flex.GCaMP6s | Syn.Flex.GCaMP6s.WPRE.SV40 | AAV1 | Addgene | 100845 |
| syn.Flex.GCaMP6m | Syn.Flex.GCaMP6m.WPRE.SV40 | AAV1 | Addgene | 100838 |
| CamKII0.4.Cre.SV40 | CamKII 0.4.Cre.SV40 | AAV1 or 9 | Addgene | 105558 |

*At the IEECR/University of Bonn:* Mice (∼6 wks of age) received ketoprofen (Gabrilen, Mibe; 5 mg/kg body weight; injection vol. 0.1 ml/10 g b.w., s.c.) for analgesia and anti-inflammatory treatment 30 minutes prior to induction of anesthesia. Then, mice were anesthetized with 2-3% isoflurane in an oxygen/air mixture (25/75%) and then placed in a stereotactic frame. Eyes were covered with eye-ointment (Bepanthen, Bayer) to prevent drying and body temperature was maintained at 37°C using a regulated heating plate (TCAT-2LV, Physitemp) and a rectal thermal probe. After hair removal and superficial disinfection, a drop of 10% lidocaine was used to locally anesthetise the area. After 3-5 mins, a flap of skin was removed about 1 cm² around the middle of the skull. Residual soft tissue was then removed from the skull with a scraper and 3% H2O2/NaCl solution. After complete drying, cranial sutures served as landmarks for the determination of injection sites. For virus injection, a burr hole was carefully drilled through the skull using a dental drill, avoiding excessive heating and injury to the meninges by intermittent cooling with sterile PBS. Coordinates were: for CA1, anterioposterior [AP] measured from bregma 1.9 mm, lateral [L] specified from midline 1.6 mm, dorsoventral [DV] from surface of the skull 1.6 mm; for dentate gyrus, AP 2.4 mm; L 1.6 mm; x3 injections at DV 2.7, 2.5, 2.1 mm. Virus particles (see **Table 1**) were slowly injected (20-100 nl/min). To prevent reflux of the injected fluid upon cannula retraction, it was left in place until 5 min post-injection, and then carefully lifted.

*At the DZNE:* A more detailed procedure was described previously (Fuhrmann *et al*. 2015; Poll *et al*. 2020). Briefly, mice (6-78 wks) were anesthetized with i.p. injection of Ketamine (0.13 mg/g) and Xylazine (0.01_Jmg/g), head-fixed using a head holder (MA-6N, Narishige, Tokyo, Japan) and placed into a motorized stereotactic frame (Luigs-Neumann, Ratingen, Germany). Body temperature was constantly controlled by a self-regulating heating pad (Fine Science Tools, Heidelberg, Germany). After skin incision and removal of the pericranium, position of the injection 34_JG cannula was determined in relation to bregma. A 0.5 mm hole was drilled through the skull (Ideal micro drill, World Precision Instruments, Berlin, Germany). Stereotactic coordinates were taken from Franklin and Paxinos, 2008 (The Mouse Brain in Stereotaxic Coordinates, Third Edition, Academic Press). Virus (see **Table 1**) was injected in two loci with the following CA1 coordinates: AP 1.95 mm; L 1.5 mm; DV 1.15 mm at a speed of 100nL/min.

*At Columbia University:* A more detailed procedure was described previously (Wenzel *et al*. 2017, 2019a). Briefly, mice (8-20 wks) were anesthetized with isoflurane (initial dose 2–3% partial pressure in air, then reduction to 1–1.5%). For viral injections, a small cranial aperture was established using a dental drill above somatosensory cortex (coordinates from bregma: AP 2.5 mm, L 0.24 mm, DV 0.2 mm), or V1 (coordinates from lambda: AP =2.5 mm, L = 0.02 mm, DV 0.2-0.3 mm), or the hippocampus (coordinates from bregma, CA1: -1.9mm, -1.6mm, - 1.6mm, CA3: -2.2 mm, -2.3 mm, -2.7 mm). A glass capillary pulled to a sharp micropipette was advanced with the stereotaxic instrument, and virus particles (see **Table 1**) were injected into putative layer 2/3 of neocortex over a 5 min period at 50nl/min, or hippocampus over 12.5 min using a UMP3 microsyringe pump (World Precision Instruments).

At *University of Zurich:* A more detailed procedure was described previously (Rupprecht *et al*. 2023). Briefly, mice (18 wks) were anesthetized using isoflurane (5% in O_2_ for induction, 1-2% for maintenance during surgery) and provided with analgesia (Metacam 5 mg/kg bodyweight, s.c.). Body temperature was maintained at 35-37°C using a heating pad. An incision was made into the skin after local application of lidocaine. Viral particles (see **Table 1**) were injected into CA1 (coordinates AP -2.0 mm, ML -1.5 mm from Bregma, DV -1.3) using a glass pipette with a manually driven syringe at a rate of approximately 50 nl/min. The injection pipette was left in place for further 5mins before being slowly retracted.

#### In vivo imaging window implantation procedure

In vivo imaging window implantation procedure: Cranial window surgery was performed to allow imaging from the dorsal hippocampal CA1/CA3 region or neocortex.

*At the IEECR, University of Bonn:* 30 minutes prior to induction of anesthesia, buprenorphine was administered for analgesia (Buprenovet, Bayer; 0.05 mg/kg body weight; injection vol.0.1 ml/20 g b.w., i.p.). Further, dexamethasone (Dexa, Jenapharm; 0.1mg/20g b.w.; injection vol. 0.1 ml/20 g b.w., i.p.), and ketoprofen (Gabrilen, Mibe; 5 mg/kg b.w.; injection vol. 0.1 ml/10 g b.w., s.c.) were applied to counteract inflammation, swelling and pain. Mice were anesthetized with 2-3% isoflurane in an oxygen/air mixture (25/75%) and then placed in a stereotactic frame. Eyes were covered with eye-ointment (Bepanthen, Bayer), body temperature was maintained at 37°C by closed-loop regulation through a warming pad (TCAT-2LV, Physitemp) and a rectal thermal probe. Throughout the course of the surgical procedure, the isoflurane dose was successively reduced to about 1-1.5% at a gas flow of ∼0.5 ml/minute. A circular craniotomy (Ø ∼3 mm) was established above the right hemisphere/hippocampus within the central opening (Ø ∼7mm) of the head plate using a dental drill. Cortical tissue was carefully aspirated until the alveolar fibers above CA1 could be visually identified. A custom-made silicon cone (top Ø 3 mm, bottom Ø 2 mm, depth 1.5 mm, RTV 615, Momentive) attached to a cover glass (Ø 5 mm, thickness 0.17 mm) was inserted and fixed with dental cement around the edges of the cover glass (see Masala *et al*. 2022). Postoperatively, all mice received analgetic treatment by administration of buprenorphine twice daily (Buprenovet, Bayer; 0.05 mg/kg b.w.; injection vol. 0.1 ml/20 g b.w., i.p) and ketoprofen once daily (Gabrilen, Mibe; 5 mg/kg b.w.; injection vol. 0.1 ml/10 g b.w., s.c.) for 3 consecutive days post-surgery. Throughout this time, animals were carefully monitored twice daily. Animals typically recovered from surgery within 24-48 hours, showing normal activity and no signs of pain or distress.

*At the DZNE*, Prior to surgery mice were anesthetized with an intraperitoneal injection of ketamine/xylazine (0.13/0.01_Jmg_Jper_Jg of body weight). Additionally, an anti-inflammatory (dexamethasone, 0.2_Jmg_Jper_Jkg) and an analgesic drug (buprenorphine hydrochloride, 0.05_Jmg_Jper_Jkg; Temgesic, Reckitt Benckiser Healthcare) were subcutaneously administered. A cranial window (Ø 3 mm) was implanted above the right hippocampus as previously described (Poll *et al*. 2020).

*At Columbia University:* For neocortical imaging, directly following virus injection, the craniotomy was covered with a thin glass cover slip (3×3mm, No. 0, Warner Instruments), which was fixed in place with a slim meniscus of silicon around the edge of the glass cover and finally cemented on the skull using small amounts of dental cement around the edge. For hippocampal imaging, a small area of cortex (around 1.5×1.5mm) above left CA1 was removed by gentle suction down to the external capsule, as described previously (Dombeck *et al*. 2010; Wenzel *et al*. 2019b). The site was repeatedly rinsed with sterile saline until no further bleeding could be observed. Then, a small UV-sterilized miniature glass plug (1.5×1.5mm, BK7 glass, obtained from BMV Optical), glued to the center of a thin glass coverslip (3×3mm, No. 0, Warner Instruments) with UV-sensitive glue, was carefully lowered onto the external capsule, and until the edges of the attached glass cover touched the skull surrounding the craniotomy. Finally, the plug was fixed in place with a slim meniscus of silicon around the edge of the glass cover and by applying small amounts of dental cement around the edge of the glass cover.

*At University of Zurich:* A more detailed procedure was described previously (Rupprecht *et al*. 2023). Briefly, 2 weeks after virus injection, the hippocampal window was implanted. Two layers of light-curing adhesive (iBond Total Etch, Kulzer) were applied to the exposed skull, followed by a ring of dental cement (Charisma, Kulzer). A 3-mm diameter ring was drilled into the skull, centered at the previous injection site. The cortex in the exposed region was carefully aspirated using a vacuum pump until the stripes of the corpus callosum became visible. The corpus callosum was left intact. A cylindrical metal cannula (diameter 3 mm, height 1.2-1.3 mm) attached with dental cement to a coverslip (diameter 3 mm) was carefully inserted into the cavity. The hippocampal window was fixed in place using UV-curable dental cement (Tetric EvoFlow, Ivoclar).

#### Two-photon Ca^2+^ imaging

A variety of standard commercially available two-photon systems were used at the different institutes to record the Ca^2+^ micro-waves.

*At the IEECR, University of Bonn*: a commercially available two-photon microscope was used (A1 MP, Nikon), equipped with a 16x water-immersion objective (N.A.=0.8, WD=3 mm, CFI75 LWD 16X W, Nikon), and controlled by NIS-Elements software (Nikon). GCaMP6s was excited at 940 nm using a Ti:Sapphire laser system (∼60 fs laser pulse width; Chameleon Vision-S, Coherent). Emitted photons were collected using gated GaAsP photomultipliers (H11706-40, Hamamatsu). Several individual tif series were recorded by resonant scanning at a frame rate of 15 Hz for a total 20-40 minutes per imaging session.

*At the DZNE*: Recordings of Ca^2+^-changes were performed with a galvo-resonant scanner (Thorlabs, Newton, USA) on a two-photon microscope equipped with a 16x water immersion objective with a numerical aperture of 0.8 (N16XLWD-PF, Nikon, Düsseldorf, Germany) and a titanium sapphire (Ti:Sa) 80 MHz Cameleon Ultra II two-photon laser (Coherent, Dieburg, Germany) that was tuned to 920 nm for GCaMP6m fluorescence excitation. GCaMP6m fluorescence emission was detected using a band-pass filter (525/50 nm, AHF, Tübingen, Germany) and a GaAsP PMT (Thorlabs, Newton, USA). ThorImageLS software (Thorlabs, version 2.1) was used to control image acquisition. Image series (896 x 480 pixels, 0.715 µm/pixel, or 640 x 256 pixels) were acquired at 30.3 Hz or 32.3 Hz.

*At Columbia University*: Neural population activity was recorded using a commercially available two-photon microscope (Bruker; Billerica, MA) and a Ti:Sapphire laser (Chameleon Ultra II; Coherent) at 940 nm through a 25x objective (Olympus, water immersion, N.A. 1.05). Resonant galvanometer scanning and image acquisition (frame rate 30.206 fps, 512×512 pixels) were controlled by Prairie View Imaging software.

*At University of Zurich*: Neuronal population activity was recorded using a custom-built two-photon microscope (see Rupprecht *et al*. 2023). Briefly light from a femtosecond-pulsed laser (MaiTai, Spectra physics; tuned to 960 nm; power below objective 40-45 mW) was used to scan the sample below a 16x objective (Nikon, water immersion, NA 0.8). Image acquisition and scanning (frame rate 30.88Hz, 622×512 pixels) were controlled by custom-written software (Chen *et al*. 2016).

#### Analysis of aberrant Ca^2+^ micro-waves

To remove motion artifacts, recorded movies were registered using a Lucas–Kanade model (Greenberg & Kerr 2009), or the ImageJ plugin moco (available through the Yuste web page, or https://github.com/NTCColumbia/moco) (Dubbs *et al*. 2016) or in the case of R-CaMP the NoRMCorre algorithm (Pnevmatikakis & Giovannucci 2017).

We determined the diameter of the calcium waves in a semi-automated fashion from the raw tif-series. Using ImageJ software, we first drew an orthogonal line across the largest aspect of each calcium wave progressing through the field of view, which resulted in a fluorescent histogram for each wave. Using custom code (MATLAB R2020b), we further analysed all histograms for each mouse and imaging timepoint. First, we applied a gentle smoothing, max-normalized each histogram, and max-aligned all histograms of a given imaging session. Then, after excluding the 10% lowest fluorescent values, the width of each calcium wave and a mean value was calculated for each time point/imaging session. Finally, the resulting pixel values were converted to micrometer, based on the respective objective (@ 512×512pxl and 1x Zoom: Nikon 16x, NA 0.8, 3mm WD: 1.579 µm/pxl; Olympus 25x, NA 1.05 2mm WD: 0.92 µm/pxl). The speed of the Ca^2+^ micro-waves was calculated from the duration and path length of the events visually identified and manually tracked in the FOV.

#### Histochemistry

To verify successful viral transduction and window position, animals were deeply anesthetized with ketamine (80 mg/kg body weight) and xylazine (15 mg/kg body weight) at the end of the respective experiment. After confirming a sufficient depth of anesthesia, mice were heart-perfused with cold phosphate buffered saline (PBS) followed by 4% paraformaldehyde (PFA) in PBS. Animals were decapitated and the brain removed and stored in 4% PFA in PBS solution for 24 hours. Fifty to 100 µm thick coronal slices of the hippocampus were cut on a vibratome (Leica). For nuclear staining, brain slices were kept for 10 min in a 1:1000 DAPI solution at room temperature. Brain slices were mounted and the green and blue channel successively imaged under an epi-fluorescence or spinning disc microscope (Visitron VisiScope).

## Acknowledgements

We sincerely thank Lea Adenauer, Laura Kück and Olga Zabashta for excellent technical support with animal husbandry and immunohistochemistry. We acknowledge the support of the Imaging Core Facility of the Bonn Technology Campus Life Sciences funded by the Deutsche Forschungsgemeinschaft (DFG, German Research Foundation) – Projektnummer 388169927.

## Funding

The work was supported by the Deutsche Forschungsgemeinschaft (DFG, German Research Foundation) with a Research Group FOR-2715 to T.K. and H.B, SPP2395 to MF and H.B, and SFB1089 to M.F, M.W. and H.B. The work was further supported by the BONFOR Program UoB (M.W.: #2019-2-04), the Hertie Network of Excellence in Clinical Neuroscience (M.W.: #P1200008), European Research Council (M.W.: StG #101039945; F.H.: AdvG #670757, M.F.: CoG#865618), the Swiss National Science Foundation (project grant 310030B_170269 and Sinergia grant CRSII5 180316 to F.H.; Ambizione grant PZ00P3_209114 to P.R.). The work was further supported by the iBehave network (M.F., H.B., M.W.). Additional support was provided by the National Science Foundation (NSF GFRP to D.A.O; 2203119 to R.Y.), National Institute of Neurological Disorders and Stroke (F99NS134209 to D.A.O. and RM1NS132981 to R.Y.), National Institute of Mental Health (R01MH115900 to R.Y.), NSF (2203119 to R.Y.), and a Vannevar Bush Faculty Fellowship (ONR N000142012828 to R.Y.)

The funders had no role in study design, data collection and interpretation, or the decision to submit the work for publication.

## Author Contributions

M.W. and T.K. conceived and planned the study. N.M., M.W. M.M, E.A., D.A.O, F.D., P.R performed experiments. F.H., R.Y., M.F., H.B. provided infrastructure and experimental expertise. M.W. and T.K. analysed data and wrote the manuscript with input from all authors.

## Conflict of Interest

None of the authors have a conflict of interest.

## Notes

### Competing Interest Statement

The authors have declared no competing interest.

### Summary of Updates

In the revised the manuscript, we changed the title to avoid any confusion with adeno viruses, included new experiments and data analyses as well as updates to both the figures and the text according to the suggestions of the referees. Our major changes include: - Additional 3 animals with conditional GCaMP6 expression supporting the use of this viral transduction procedure as an alternative to avoid the Ca2+ micro-waves - A new analysis of age-related changes and statistical analysis between different conditions - A revised manuscript text including discussion points suggested by the reviewers - An emphasis on the GECI expression level and cell density as the major factors, with the CA1/CA3 regions being particularly prone to generate Ca2+ micro-waves following synapsin-dependent expression.

