## Supplementary Material Legends for "Aberrant hippocampal Ca^2+^ micro-waves following synapsin-dependent adeno-associated viral expression of Ca^2+^ indicators"

**Video 1.**

GCaMP6s two-photon calcium imaging in hippocampal CA1 region, around 100 µm beneath hippocampal surface (stratum pyramidale), ~4 weeks after transduction of AAV1 particles containing pAAV.Syn.GCaMP6s.WPRE.SV40 (Addgene plasmid #100843) in a mature bl6 wildtype mouse. Imaging wave length = 940 nm, acquisition speed = 15 frames/sec. Movie played at 5x acquisition speed. Imaging performed at IEECR/University of Bonn.

**Video 2.**

GCaMP6s two-photon calcium imaging in hippocampal CA1 region, around 100 µm beneath hippocampal surface (stratum pyramidale), ~2 weeks after transduction of AAV1 particles containing pAAV.Syn.GCaMP6s.WPRE.SV40 (Addgene plasmid #100843) in a ~2 months old transgenic mouse model of genetic epilepsy (heterozygous *Scn2a*^A263V^ mouse). Imaging wave length = 940 nm, acquisition speed = 15 frames/sec. Movie played at 5x acquisition speed. Imaging performed at IEECR/University of Bonn.

**Video 3.**

R-CaMP1.07 two photon calcium imaging in hippocampal CA1 region, around 100µm beneath hippocampal surface (stratum pyramidale) ~10 weeks after transduction of AAV1 particles containing ssAAV-9/2-hSyn1-chI-RCaMP1.07-WPRE-SV40p(A) (Viral Vector Core UZH #V224-9) in a mature (~5 months) bl6 wildtype mouse. Imaging wave length = 960 nm, acquisition speed = 30.88 frames/sec. Movie played at 5x acquisition speed. Imaging performed at Neuroscience Center Zurich (UZH).

**Video 4.**

GCaMP6s two-photon calcium imaging in hippocampal CA1 region, around 100 µm beneath hippocampal surface (stratum pyramidale), ~7 weeks after transduction of AAV1 particles containing pAAV.Syn.GCaMP6s.WPRE.SV40 (Addgene plasmid #100843) in a ~3 months old transgenic mouse (same as in suppl. video 2; *Scn2a*^A263V^ model of genetic epilepsy). Imaging wave length = 940 nm, acquisition speed = 15 frames/sec. Movie played at 5x acquisition speed. Imaging performed at IEECR/University of Bonn.

**Video 5.**

Same animal (*Scn2a*^A263V^ model of genetic epilepsy) and timepoint of imaging as in suppl. video 4. GCaMP6s two-photon calcium imaging in hippocampal CA1 region, around 25 µm beneath hippocampal surface (stratum oriens), ~7 weeks after transduction of AAV1 particles containing pAAV.Syn.GCaMP6s.WPRE.SV40 (Addgene plasmid #100843). Imaging wave length = 940 nm, acquisition speed = 15 frames/sec. Movie played at 5x acquisition speed. Imaging performed at IEECR/University of Bonn.

**Video 6.**

GCaMP6s two-photon calcium imaging in hippocampal CA3 region, stratum pyramidale, ~7 weeks after transduction of AAV1 particles containing pAAV.Syn.GCaMP6s.WPRE.SV40 (Addgene plasmid #100843) in a mature bl6 wildtype mouse. Imaging wave length = 940 nm, acquisition speed = 30.206 frames/sec. Movie played at 5x acquisition speed. Imaging performed at Columbia University.
